# Gel-Shrink Softlithography: Enabling 3D Tissue Culture Microstructure Miniaturization and Tacky Super-Soft Material Molding

**DOI:** 10.1101/2023.09.27.559860

**Authors:** Cheng-Kuang Huang, Chwee Teck Lim

## Abstract

In this study, we introduce an innovative hydrogel soft lithography technique that leverages the significant volume change of hydrogels during hydration. Through a systematic desiccation process applied to hydrogels containing microfeatures, we achieved a remarkable 2-fold reduction in feature dimensions. These shrunken hydrogel structures were then readily molded using standard soft lithography methods. Furthermore, we demonstrated the technique’s utility in rectifying surface imperfections commonly found in 3D printed structures, offering promising prospects in fields like tissue engineering. Additionally, our study discovered a unique capability of hydrated hydrogel molds in molding super-soft, tacky elastomeric materials, exemplified by CY25-276 and low cross-linker ratio PDMS. This development opens doors to studying cellular forces through traction force microscopy to a broader spectrum of 3D structural contexts. In support of our efforts in microfabrication advancement, we explored multicellular responses to surface curvatures of diminishing dimensions. Our findings reveal a curvature-dependent alignment pattern: cells residing on convex surfaces align with the direction of least curvature, while those in concave regions align with the direction of the strongest curvature. These results underscore the critical influence of geometric complexity in biological systems and highlight the need for continuously improving methods to replicate geometric qualities relevant to these environments.

## INTRODUCTION

Surface curvature stands as a foundational mathematical concept used to characterize the geometrical attributes of objects. When this concept is applied to biological structures, it offers a quantitative framework to precisely delineate the intricate geometry of various components within organisms: from the spherical configurations of organs like the eyes and alveolar acini to the cylindrical formations exhibited by arteries and intestinal villi, among others. Notably, these surface curvatures are intrinsically linked to the underlying biological functions *(1)*.

We posit that cells, as the fundamental building blocks of life, have evolved mechanisms for detecting and responding to curvature cues within their microenvironment. The significance of this curvature-sensing ability extends beyond embryonic development and morphogenesis, encompassing the vital processes of tissue maintenance and repair in fully developed organisms, which are essential for homeostasis.

The empirical evidence supporting this hypothesis is substantial, revealing distinct cellular responses to a diverse array of cellular-scale (10^1^-10^3^ μm) curved features *(2, 3)*, ranging from spherical *(4, 5)* and cylindrical *(6–8)* to toroidal *(9)* and wavy *(10–12)* surfaces. The observed curvature-induced effects encompass a wide spectrum of phenomena, including changes in cellular morphology, preferences in cytoskeletal and nuclei alignment, nuclei deformation, guided migration, alterations in cell differentiation, and disparities in cell proliferation rates. For instance, Hwang et al. *(8)* illustrated that L929 mouse fibroblasts exhibit a pronounced alignment preference along the long axis of narrower PLGA microfibers, thereby underscoring the role of surface curvature in directing cellular behavior. Pieuchot et al. *(12)*, conversely, demonstrated the avoidance of convex hill regions by human mesenchymal stem cells during migration along sinusoidal surfaces. More recently, Huang et al. *(4)* elucidated the phenomenon of cell cycle arrest in rat embryonic fibroblasts confined within concave spherical pores, with diminishing effects noted at smaller curvatures corresponding to larger pores.

In a recent study, our research team devised a versatile microfabrication methodology designed to generate hemi-cylindrical wave structures *(13)*. These structures feature half-period dimensions spanning from conventional cell sizes to dimensions exceeding several hundred microns. Notably, our approach distinguishes itself from extant methodologies for producing wavy patterns in that it yields developable surfaces alternating between convex and concave features characterized by similar degrees of curvature. This symmetrical dimensionality is of paramount significance, as it eradicates the intrinsic ambiguity encountered in studies featuring stark curvature disparities between convex and concave regions. In particular, the ability to discern between the influence of curvature magnitude and the direction of curvature (convex or concave) is preserved. Furthermore, our novel wave substrates exhibit an “edge-free” attribute, wherein convex regions transition seamlessly into concave regions, and vice versa—a property referred to as “edge-free” in the relevant literature *(12)*. This characteristic is highly advantageous, as it mitigates the geometric complexity associated with structures featuring intervening edges and flat regions, thus minimizing the potential for compounded effects.

Building upon our established microfabrication methodology, this study endeavors to expand our capacity to produce curved substrates with increasingly diminutive feature dimensions. In previous investigations, we had relied upon commercial glass rods to fashion hemi-cylindrical wave substrates. However, our endeavors were confined to dimensions commencing at 100 μm, as smaller rods were not commercially available. To surmount this limitation, we resorted to an ad hoc solution, involving the stretching of soft elastomeric molds, to reduce lateral dimensions. While this approach yielded satisfactory results for developable surfaces such as cylindrical waves, it could not be applied effectively to diminish the dimensions of isotropic features. For example, stretching spherical substrates led to the formation of slender ellipsoidal structures, deviating from the objective of producing smaller spherical ones.

In the present work, we introduce a novel hydrogel soft-lithography technique designed to afford the capacity to reduce the dimensions of various surface microfeatures in a continuous manner. With this innovative lithography approach, we have successfully reduced the dimensions of hemi-cylindrical waves to the scale of a single cell width (∼25 μm). Furthermore, we have demonstrated the adaptability of our method in adjusting the dimensions of isotropic round well features. Additionally, our methodology proved instrumental in rectifying surface defects inherent in 3D printed surfaces, a common challenge encountered in biomaterials engineering studies relying on techniques such as Two-photon lithography to create microstructures *(14)*. These periodic surface defects can introduce spurious responses in tissues cultivated on 3D printed materials, as cells have readily been found to respond to even nanometric topography *(15)*. This underscores the practical utility of our approach in enhancing precision and reliability within such investigative domains.

### GEL-SHRINK SOFTLITHOGRAPHY

Our Gel-Shrink softlithography method capitalizes on the unique properties of hydrogels, specifically their ability to significantly contract when water is removed. We initiate the process by molding polyacrylamide (PAM) against microstructured substrates. Controlled desiccation gradually eliminates water from these hydrogel replicas, resulting in rigid molds. These molds are then used to shape silicone elastomer, yielding structures that are approximately half the size. This procedure can be repeated for further size reduction, offering a highly scalable and precise microfabrication approach. The steps of our procedures are as follows (schematically shown in Figure 1):

**Figure 1.**
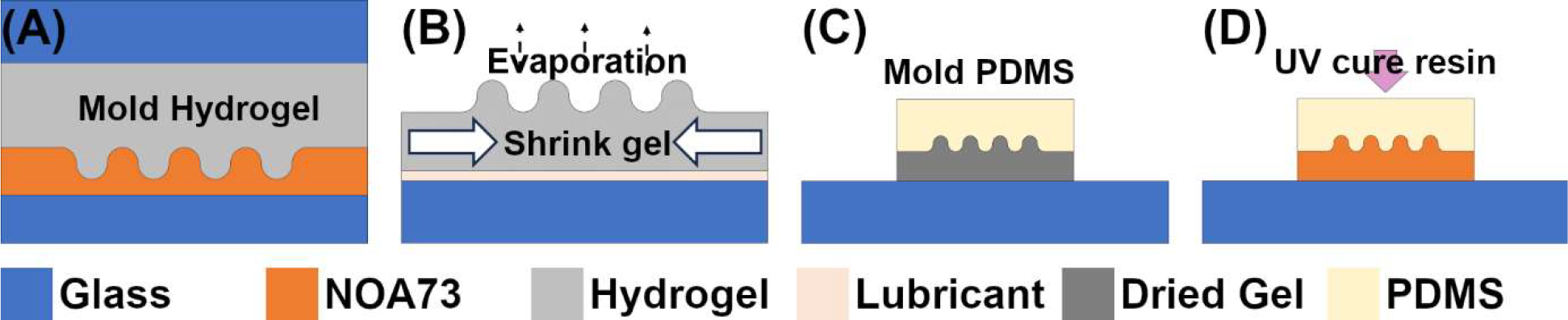
Schematic illustration of Gel-Shrink softlithography. (A) step 1, mold hydrogel against micropattern. (B) step 2, shrink hydrogel on lubricated surface through gradual evaporation. (C) step 3, clone the dried gel with silicones. (D) step 4, create a robust replica of the silicone clone using UV curable resin. The dimensions of the micropatterns can be further reduced through recursive iteration of the whole procedure.

1. *Polyacrylamide Preparation and Molding (Figure 1a)* In the first step, we prepared a polyacrylamide precursor solution by combining specific components: 200 μl of 40% polyacrylamide solution (Bio-Rad), 200 μl of 2% bisacrylamide solution (Bio-Rad), 580 μl of MilliQ water, and 1.5 μl of TEMED (Bio-Rad). After adding 2 μl of 10% APS (Sigma-Aldrich) and mixing, we applied the mixture to molds with microstructure features. A cover glass was placed over the mold, and the polyacrylamide was allowed to gel for 30 minutes. Then, we gently removed the polyacrylamide from the mold, lifted it from the supporting glass, and immersed it in MilliQ water.
2. *Controlled PAM Gel Drying (Figure 1b)* In the next step, we focused on controlled desiccation of the PAM (polyacrylamide) gel slab to minimize deformation. We smeared the supporting surface with PDMS (Sylgard 184, Dow Inc., US) monomer liquid Part A to facilitate smooth drying. After placing a PAM slab on this surface, we wicked away excess water using lint-free: we maintained a lint- and dust-free environment to avoid particulate on the surface, which would induce deformations. Finally, we slowed down evaporation by covering the slab with a lid, typically from a 35mm culture dish. If one wishes to speed up the desiccation process, the hydrogel slabs can be placed in ethanol for 1 hour first for a pre-dehydration.
3. *Molding Silicone Elastomer on the Dried Gel (Figure 1c)* Moving on to the third step, the PAM gel slabs typically fully dried into transparent, brittle plastic-like sheets within 24-48 hours, contingent upon environmental humidity levels. It’s important to note that some deformations might occur at the slab edges due to variations in the rate of water loss. However, these deformations generally do not impact the patterns located in the central regions of the slabs. If further reduction of deformations is desired, the desiccation process can be intentionally extended.
4. *Creating a robust resin mold and repeating to further reduce dimensions (Figure 1d)* In the fourth step of our process, we start with the PDMS (polydimethylsiloxane) clone of the PAM (polyacrylamide) template and utilizing Norland Optical Adhesive 73 ((NOA73, Norland Products Inc., UK)) to create a robust replica of the PAM pattern from the PDMS clone. In brief, a drop of NOA73 is applied to the PDMS mold, pressed against a glass slide, and cured using a UV-KUB 9 (KLOE, France) for approximately 2 minutes at 25% power. This curing process solidifies the NOA73 resin, creating a robust mold adhered to the glass. The mold was then baked overnight to enhance its adhesion to glass. We can achieve further reductions in feature dimensions through repeating the above steps with this new resin mold.

## RESULTS AND DISCUSSIONS

Figure 2a shows a visual comparison between a desiccated PAM substrate (right) and its original NOA73 100 μm (half-period width) hemicylindrical wave mold (left). While Figure 2b shows a similar comparison but for a repeated iteration; the NOA73 mold now contains the reduced ∼50 μm hemicylindrical wave features. Figure 2c-e are bright-field micrographs (plane and cross-section view) of the original 100 μm waves, the reduced ∼50 μm waves and the second time reduced ∼25 μm waves, respectively. These results show that we can effectively reduce the dimensions of our wave structures by half while maintaining their smoothness and symmetry.

**Figure 2.**
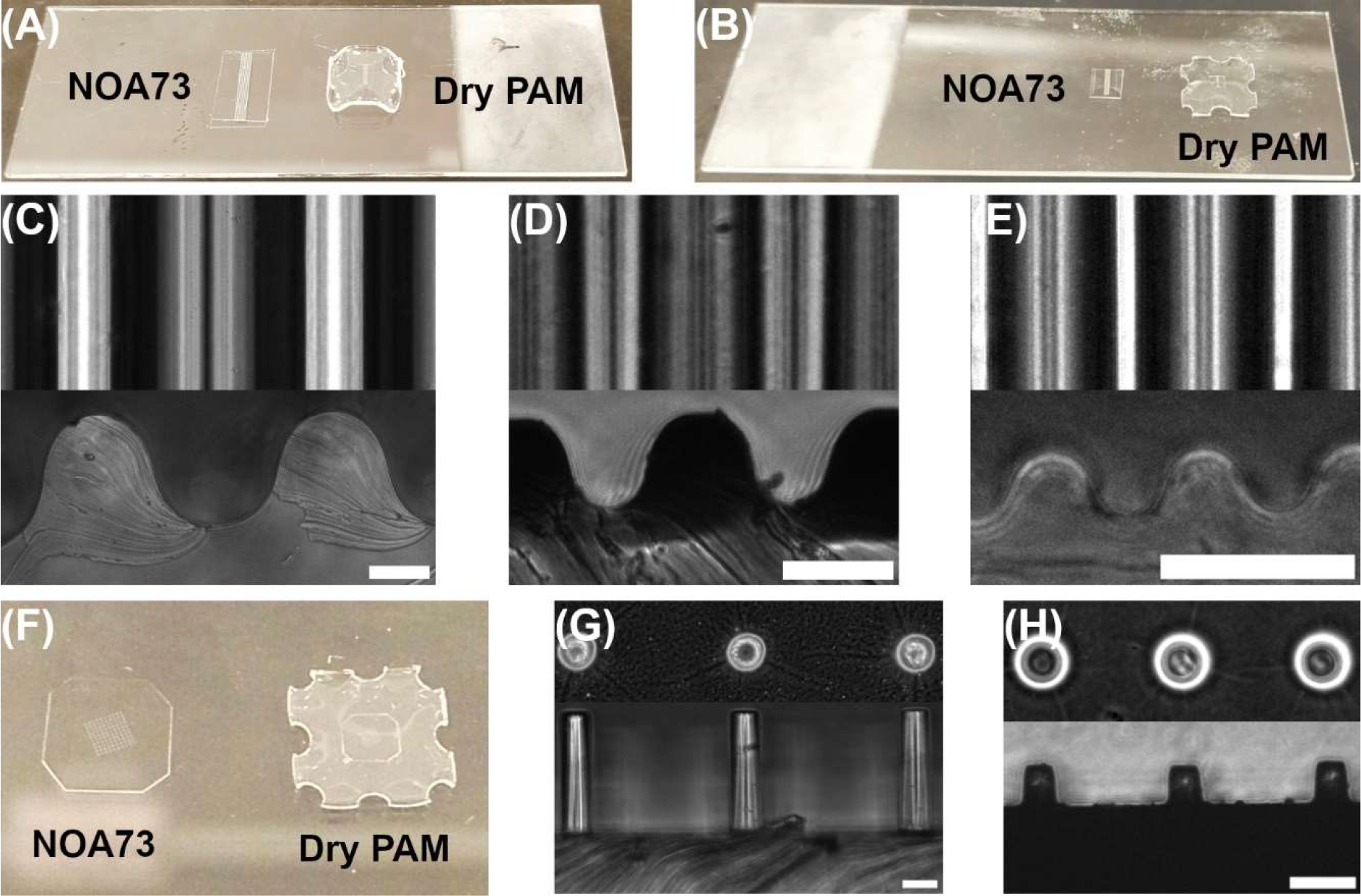
(A) a photograph showing a fully dried PAM gel with reduced wave micropatterns placed next to its original resin mold (NOA73). (B) a photograph showing a resin replica of the dried PAM in (A) and the second iteration of fully dried PAM gel with further reduced micropatterns. (C), (D), and (E) bright-field micrographs showing the plane (top) and cross-section (bottom) view of the wave microstructures with dimensions of ∼100 (original), ∼50 (1^st^ reduction), and ∼25 (2^nd^ reduction) μm, respectively. (F) a photograph showing a fully dried PAM gel with reduced circular pillar patterns next to its original resin mold. (G) and (H) bright-field micrographs of the plane (top) and cross-section (bottom) view of the circular pillar microstructures with dimensions of ∼50×200 and ∼25×50 μm, respectively. Scale-bars: 50 μm.

To show our method provide structural reduction isotropically, we applied similar treatments to ∼50 μm round pillar structures, which were originally created using standard soft-lithography techniques *(16)*. Figure 2f shows the reduced PAM substrate (right) next to the original NOA73 mold (left). Figure 2g and h are bright-field micrographs (plane and cross-section view) of the original ∼50 μm pillar structures and the reduced ∼25 μm pillar structures, respectively. The images clearly show reductions in all three dimensions.

In recent times, nano 3D printing methods like nanoscribe have garnered attention from biomaterial engineers due to their versatility in crafting customized structures and surfaces for 3D tissue culture applications *(14)*. However, these methods have inherent downsides, notably the characteristic textures generated by layer-by-layer deposition, ranging from several hundred nanometers to single microns. These textures can potentially trigger unintended cellular responses when overlaid with cells *(15)*, complicating the assessment of structure performance. To address this challenge, our method, which can reduce feature dimensions can in theory be adapted to refining surface imperfections in 3D printed structures. As a proof of concept, we molded PAM gels onto a FDM (Fused Deposition Modeling) printed slabs (Figure 3a), reduced them, and cloned accordingly. Figure 3b shows the micrograph of the original FDM surface defect pattern and Figure 3c shows the same defects after reduction. Evidently, the reduced patterns contain considerably smaller surface undulations than the original. Figure 3d shows the reduced FDM-patterned PAM substrate (right) next to the original NOA73 FDM patterned mold (left). In a specific use-case scenario, our method can be effectively applied by creating structures slightly larger than intended and subsequently reducing them to the desired dimensions. Doing so, one can obtain smoother surfaces for the intended microstructures compared to direct printing at the target length scale.

**Figure 3.**
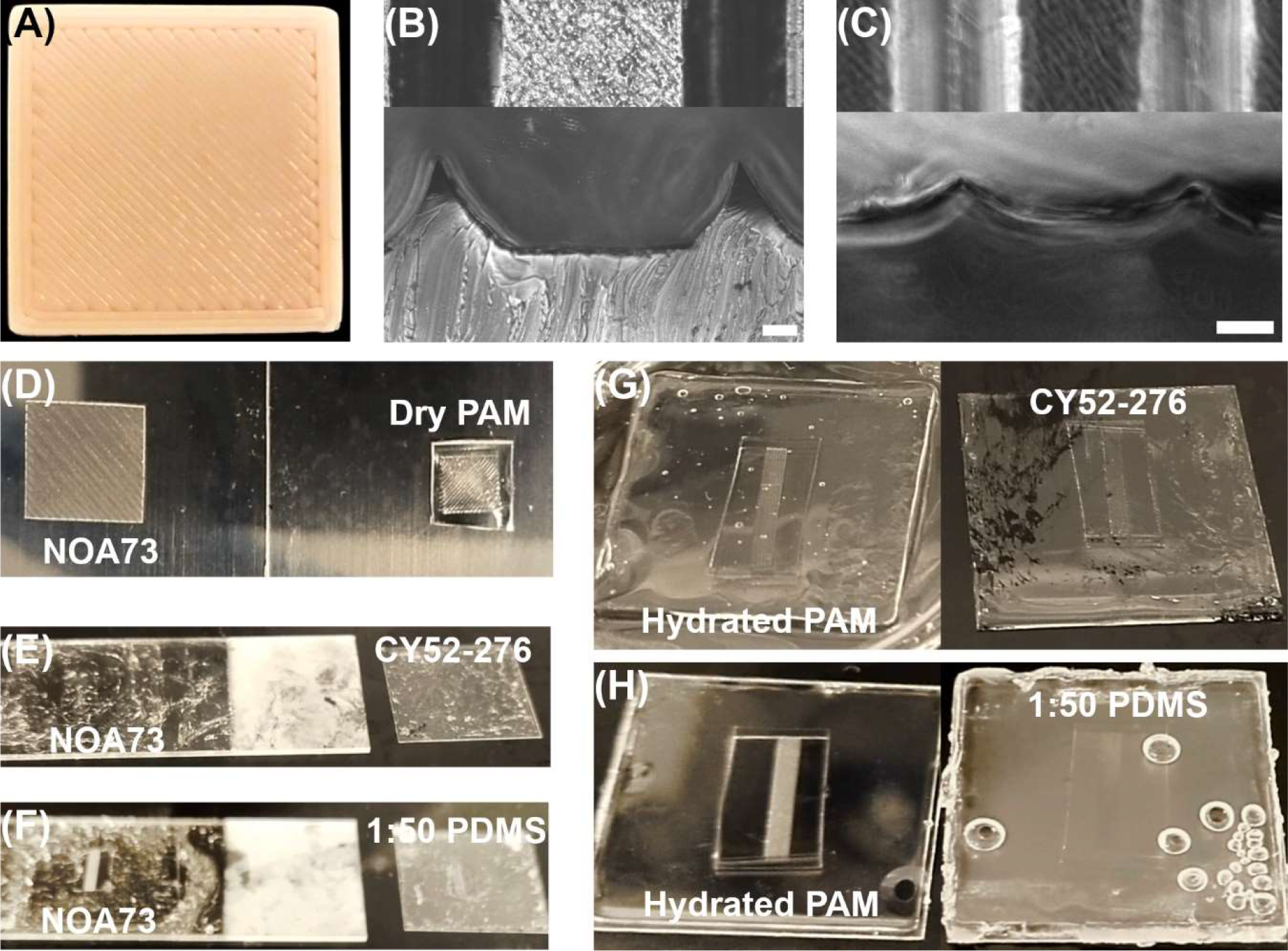
(A) a 2×2 cm PLA slab printed using a custom-built Kossel FDM printer. (B) bright-field micrographs showing the plane (top) and cross-section (bottom) view of the surface defects of (A). (C) micrographs of the smoothened surface shown in (B) through our Gel-Shrink method. (D) show the fully dried PAM gel molded against the resin replica of (A). (E) and (F) show the impossibility of de-molding super-soft tacky materials from micropatterned resin molds. (G) show photographs of successfully de-molded CY52-276 from molds of hydrated hydrogels on the left. (H) show the same in (G) but for the de-molding super-soft PDMS. Scale-bars: 50 μm.

Another notable and interesting application of our microfabrication method is its ability to mold super-soft and tacky materials like CY52-276 (Dow Inc., US) silicone *(17)*, frequently employed in cellular traction force microscopy studies. The challenge in studying cellular forces lies in the requirement for materials to be sufficiently soft to be deformed by cells. However, many commonly used inorganic materials for this purpose, silicones in particular, tend to be sticky when cured. While these materials are strictly viscoelastic, researchers in the field of Traction Force Microscopy (TFM) often assume that the forces involved (typically on the order of nano newtons) are small and temporary enough for materials to operate within their linear regimes when considering cellular effects. However, when extending this method to study cellular forces on 3D structures, a significant challenge arises. The cured elastomer naturally binds to the mold, making it impossible to release. Figure 3e and f show results of attempts to mold our wave structures with CY52-276 and 1:50 (low cross-linker content) PDMS, respectively.

A recent solution involved creating sacrificial molds by solidifying sugar on molds, subsequently using it to mold super-soft elastomers *(18)*. The substrate can then be obtained by dissolving the sugar in water upon heating.

Our method, however, overcomes this binding challenge using a different mechanism. The method capitalizes on the fact that hydrogels retain water, rendering hydrophobic elastomers unable to bind to the polymer material. It’s crucial to clarify that this process is distinct from reduction microfabrication in Figure 2. Here, for the molding process to be effective, we must utilize the swollen, wet gels themselves. In brief, PAM gel substrates immersed in MilliQ water are first carefully transferred to a surface such as glass or plastic. Excess water was then removed using lint-free paper. After this, CY52-276 silicone in a 1:1 ratio is poured onto the feature side of the PAM gel, followed by gently pressing a glass cover-slip against the elastomer drop. Care must be taken to ensure that the liquid covers all edges to maintain the gel’s hydration. Once the elastomer has solidified, we can easily detach the patterned super-soft substrate from the PAM gel using a razor blade along the edges. Figures 3g and h show representative results of successfully transferring micropatterns from the hydrogel slab (left) to substrates made of CY52-276 (right) and a 1:50 (cross-linker/monomer) PDMS (right), respectively.

To demonstrate the influence of curved surfaces at cellular length scales on multicellular behavior, we cultured confluent MDCK monolayers on our hemicylindrical wave substrates for periods of 48 and 72 hours. We then assessed actin cytoskeletal alignments at these two time points across different dimensional conditions, including 200, 100, and 50 μm half-period widths, respectively. We also conducted similar experiments on rectangular wave substrates for comparison. Prior to analysis, the z-stack images were unwrapped and mapped to 2D, this should facilitate the accurate analysis of F-actin alignment downstream. Representative flattened 3D actin fluorescent micrographs of our MDCKs on both hemicylindrical and rectangular waves are presented in the image columns in Figure 4a and b, with each column displaying the two time-points. Notably, these images reveal that actin alignment tends to be more pronounced on the hills (convex regions) compared to the valleys (concave regions). Meanwhile, alignment on the rectangular waves appears to be more randomly orientated for the larger two dimensions. The images for the two time-points of the control are shown Figure 4c, which appears randomly oriented over several cell lengths.

**Figure 4.**
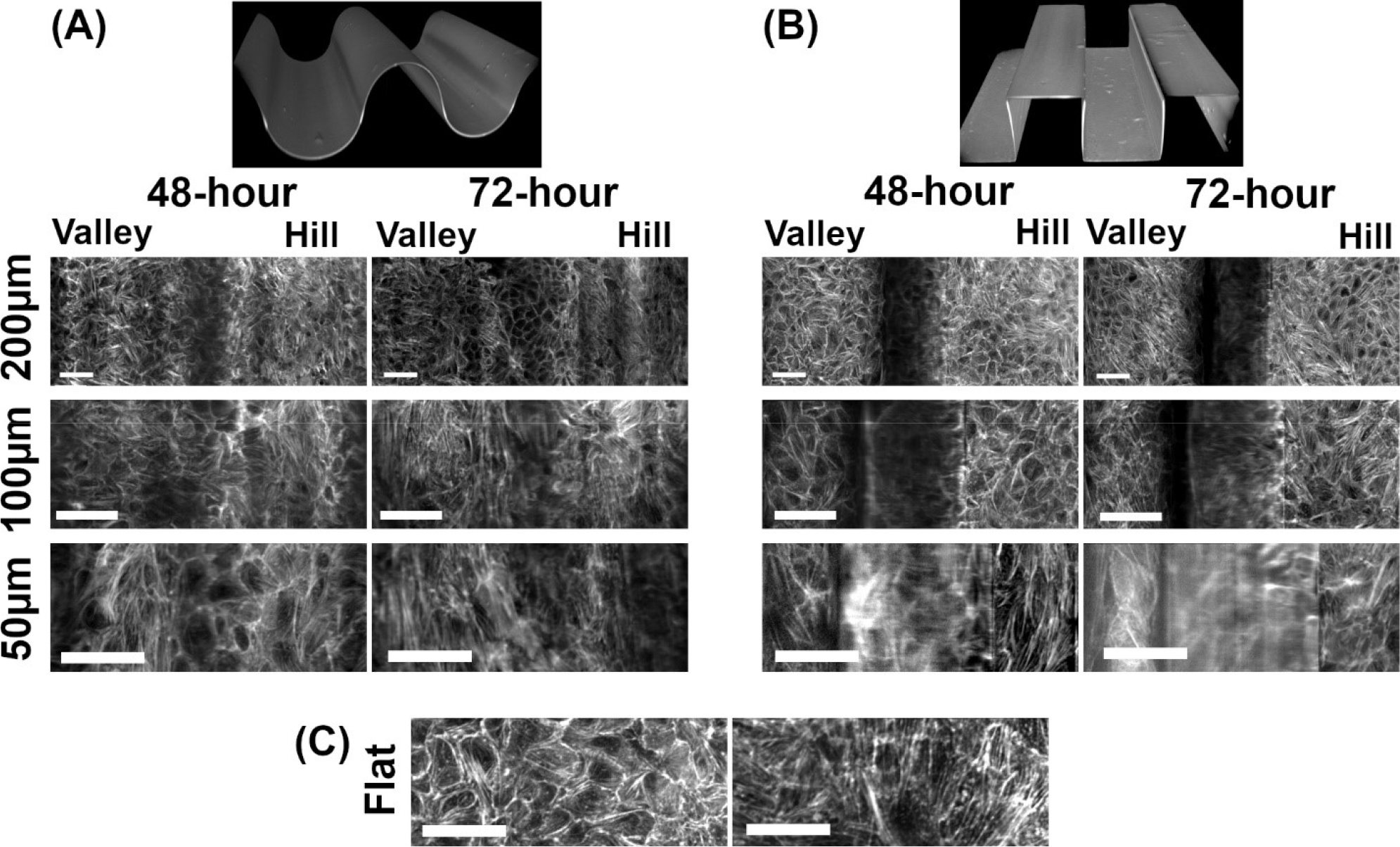
(A) 2D-mapped confocal fluorescent z-stacks showing actin distribution in MDCK monolayers cultured on hemicylindrical wave structures exemplified in the top 3D reconstruction. Conditions vary from 200-50 μm half-period width (top-down) to 48-72-hour after cell seeding. (B) the same as shown in (A) but for MDCK monolayers cultured on rectangular waves exemplified in the top 3D reconstruction image. (C) actin distribution in MDCK monolayers on planar substrates over similar time-points. Scale-bars: 50 μm.

We then conducted a detailed analysis of actin alignments of the images. The results for actin alignment on cylindrical and rectangular waves are shown in the polar-plots in the two columns of Figure 5a and b, again each with two time-points. The results confirmed that cells residing on the hills (green bars) of the hemicylindrical waves exhibit stronger alignment with the pattern in all dimensional conditions and across the two time-points. In contrast, actin alignment in the valleys (orange bars) of 100 and 200 μm conditions exhibited an intriguing anti-parallel configuration to the pattern. This anti-parallel valley actin arrangement on the 100 μm waves shifted towards alignment at the 72-hour mark while the 200 μm valley actin alignment remained anti-parallel. In the smallest 50 μm dimension (representing the strongest curvature) alignment with the pattern is found in both the valleys and hills and in both time-points.

**Figure 5.**
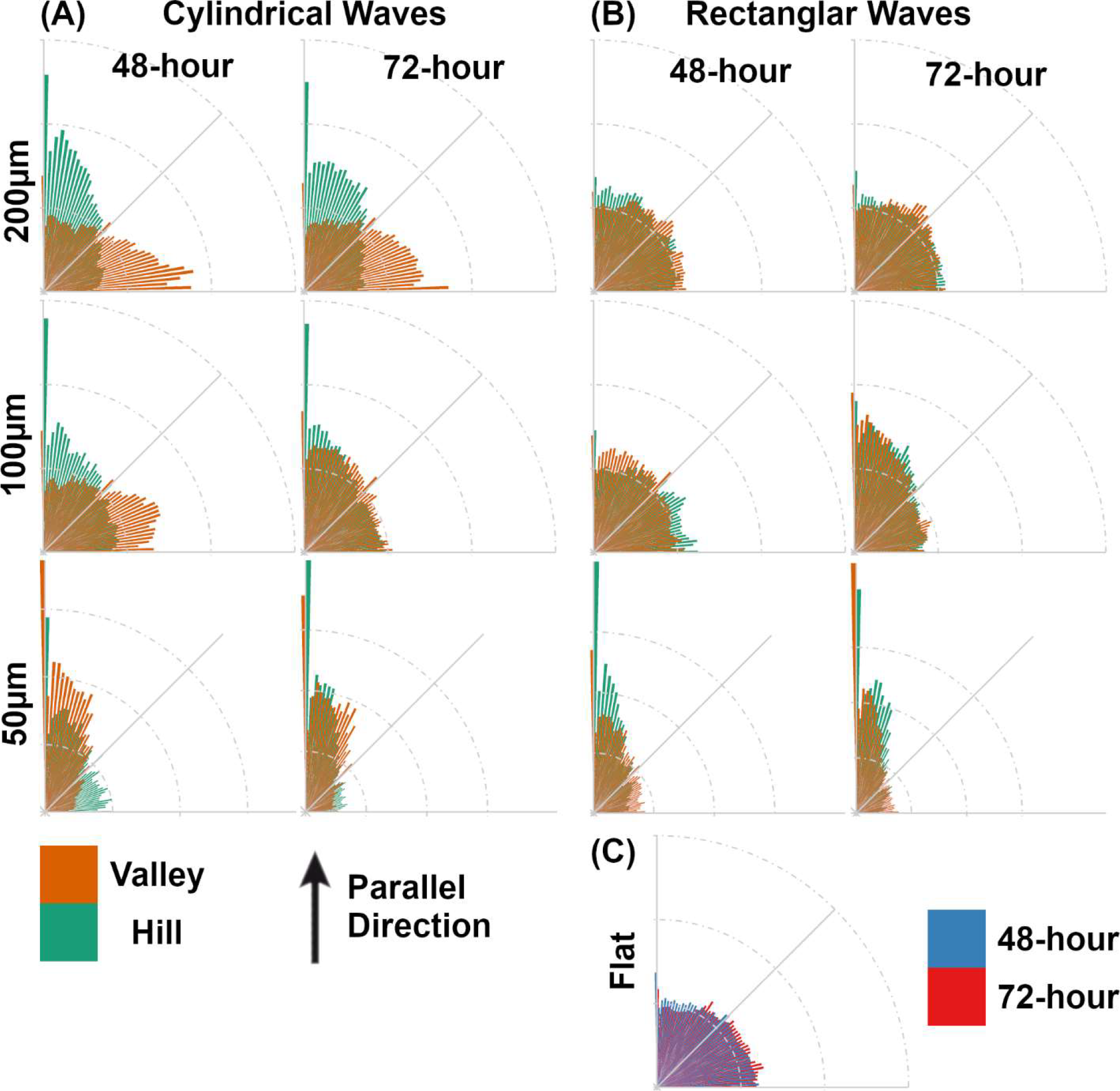
Polar plots showing actin alignment extracted from 2D-mapped fluorescent z-stacks. (A) plots for cells on the hemicylindrical waves with dimension conditions running vertically, and temporal conditions laterally. (B) is the same as in (A) but for alignment extracted from cells on rectangular patterns. (C) show the alignment for cells on planar surfaces over the two time-points.

For comparison, on the rectangular waves, a similar strong alignment was observed in the 50 μm condition. However, what distinguishes the rectangular patterns from the curved ones is the absence of alignment disparity between cells on rectangular hills and valleys of the larger two dimensions. Specifically, cellular actin alignment in both regions appeared random in the 100 and 200 μm conditions at 48 hours. This random alignment persisted in the 200 μm condition even at the 72-hour mark. An intriguing observation at this time-point was that actin alignment in both hill and valley regions aligned with the pattern on the 100 μm condition.

As a control, actin alignment on planar regions was also analyzed and shown in Figure 5c, revealing that alignment remained random at both time-points.

Subsequently, we subjected our alignment data to statistical analysis, and the distributions featured in Figure 5 were represented as boxplots in Figure 6a, illustrating results for the 48- and 72-hour time-points in the top and bottom row, respectively. It is evident that a robust alignment is consistently observed in the 50 μm conditions on the cylindrical waves at both time-points. On the other hand, for the 100 and 200 μm conditions there are clear disparities in cellular actin alignment between hill and valley regions, with hills displaying parallel alignment and the valleys showing anti-parallel alignment. Two-way ANOVA of the results revealed significance both between valley and hills (48-hour: p = 0.006, 72-hour: p = 0.007), and across the dimension conditions (48-hour: p = 7×10^-5^, 72-hour: p = 7×10^-7^). Valley-hill pairwise significances from the post hoc multiple comparisons are annotated on the boxplots.

**Figure 6.**
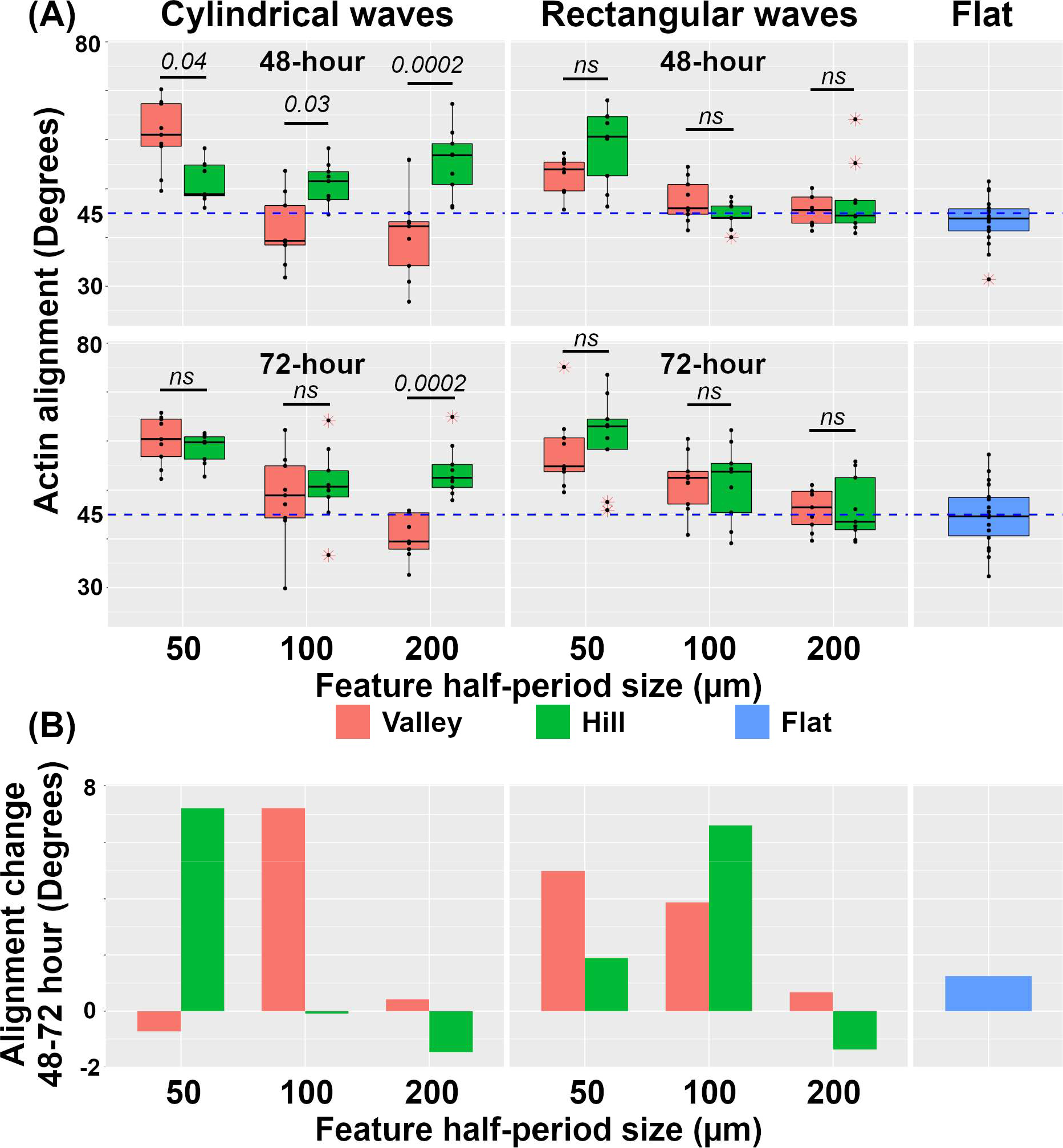
(A) show boxplots of the alignment data in Figure 5 with the top row showing data for the 48-hour mark and the bottom row the 72-hour mark. The first column shows data for cells on the hemicylindrical waves, while the second column shows data for cells on the rectangular waves. (B) is a bar chart showing the difference in actin alignment over the two time marks. The first panel for hemicylindrical waves and the second panel for the rectangular waves. The planar control is shown in the last panel.

As for the rectangular patterns, strong alignment is again noticeable in both hill and valley regions for the 50 μm patterns at both time-points. In contrast, cellular actin exhibits random arrangement on larger dimensions at the 48-hour mark. At the 72-hour time-point, actin alignment begins to manifest in both valley and hill cells in the 100 μm conditions. Two-way ANOVA of the results showed no significance between valley and hills (48-hour: p = 0.2, 72-hour: p = 0.6), but significance across the dimension conditions (48-hour: p = 7×10^-7^, 72-hour: p = 6×10^-6^). Valley-hill pairwise significance from the post hoc multiple comparisons are annotated on the boxplots.

The alignment change between the two time-points is shown in the bar-chart in Figure 6b. Overall, F-actin alignment tends to shift towards parallel alignment with the pattern over time. This temporal behavior appears to have the least effect on the largest dimension conditions in both hemicylindrical and rectangular waves. And while the effect is consistent on rectangular waves regardless of valley or hill position in the 50 and 100 μm conditions, the behavior was found to be spatially dependent in the hemicylindrical waves. Specifically, hill cells exhibit increased alignment in the 50 μm conditions, whereas such increased actin alignment happened in the valley cells in the 100 μm conditions.

Remarkably, despite the similar environmental conditions experienced by the cells in both topographical settings, a clear disagreement in actin alignment is observed between valley and hill cell-sheets in the hemicylindrical waves. Contrastingly, their counterparts on rectangular waves instead exhibit coordinated alignment in both valleys and hills. Our observations support the notion that surface curvature, and consequently geometric complexity, can finely regulate multicellular behavior, all else being equal.

It is noteworthy that the range of environmental conditions conducive to cell survival is limited, whereas the combination of surface curvatures defining tissue geometric complexity offers virtually infinite possibilities without impacting cell survival. Thus, for cells to be able to fine tune their behavior with regards to the curvatures of their environment will no doubt endow them with a greater capacity to engender diverse functions. And by virtue of this, it is not unreasonable to posit that geometrical cues could play a significant role in many biological processes, from development to homeostasis.

In conclusion, our Gel-Shrink softlithography technique introduced in this study represents a significant advancement in microfabrication methodologies. Its ability to provide precise control over feature miniaturization makes it a valuable addition to the repertoire of microfabrication tools for biomaterials engineering. The practical utility of this method extends to various crucial areas. Firstly, it effectively reduces the dimensions of microstructures, presenting promising applications in fields such as microelectronics, microfluidics, and 3D tissue-cultures where fine control over feature sizes is essential. Moreover, our method can be adapted to eliminating surface imperfections in 3D printed structures. The issue is particularly important in domains like tissue engineering and medical device fabrication, where surface quality is paramount. Our study also demonstrates the technique’s applicability in molding super-soft tacky materials. This capability holds significant implications for cellular traction force microscopy studies, facilitating the study of cellular forces on a greater variety of 3D structures. Finally, our study of multicellular structures on hemicylindrical waves reveals the profound impact of surface curvature at the cellular scale. The sensitivity of actin cytoskeletal alignments to alterations in surface curvature underscores the critical role of geometric complexity in biological systems. This understanding can help reshape our approaches to various applications, particularly in tissue engineering and regenerative medicine.

## EXPERIMENTAL SECTION

### Cell culture, staining and imaging

The Madin Darby Canine Kidney type II cells (MTOX1300, ECACC collection purchased from Sigma-Aldrich) were cultured in Minimum Essential Medium (MEM, Gibco) supplemented with 5% FBS, 1× Penicillin/streptomycin, 1× GlutaMax, 1× Sodium Pyruvate, and 1× non-essential amino acids (all from Gibco). The cell line was maintained at 37°C in 5% CO2 and sub-cultured every other day. Cells were typically disposed of before reaching 25 passages.

To facilitate cell adhesion on the PDMS wave substrates, an extracellular matrix coating was necessary. Initially, the surface was activated using oxygen plasma for 3 minutes.

Subsequently, 500 μl of a 20 mM acetic acid solution containing 50 μg of collagen-I from bovine skin (C4243, Sigma-Aldrich) was applied to the substrate, which was supported on a 22×22 mm cover-glass and allowed to incubate for 2 hours at room temperature. Following this, the collagen-I solution was removed, and the substrate was air dried for 1 hour. Coated substrates were stored at 4 °C and typically utilized within a week. Before commencing experiments, the coated substrates were re-immersed in 1× phosphate buffered saline (PBS, P4417, Sigma-Aldrich) in a 35 mm culture dish and subjected to 15 minutes of UV sterilization within a biosafety cabinet.

To investigate MDCK responses on the curved substrates, cells were initially suspended in imaging media (MEM without phenol red, with the mentioned supplements and 2% serum). Subsequently, they were seeded onto the substrates at a density of approximately 2.5×10^5^ cells/cm^2^. To achieve an even distribution of cells, a figure of eight motion was applied to the sample dish. The monolayers were then fixed for fluorescence imaging at 48- and 72-hours postseeding.

For F-actin staining, monolayers reaching desired time-points were rinsed in 1× PBS followed by fixing in 4 % formaldehyde (28906, Thermofisher Scientific) for 15 minutes. After two 1× PBS washes, the cells were permeabilized with 0.1 % Triton X-100 (X100, Sigma-Aldrich) for 3 minutes. The cells were then twice washed with 1× PBS, and stained with Alexa Fluor 647 (A30107, Thermofisher Scientific) in 1 % BSA (Sigma Aldrich) at 0.5× for 1 hour. It is imperative that while performing any fluid exchange the samples are never exposed to air, this prevents the liquid tension in the valleys from tearing the monolayers.

For imaging fixed and stained samples, they were typically elevated using spacers and positioned upside down in 1× PBS on cover-glasses. These prepared samples were then secured in a stainless-steel cover-glass holder (SC15022, Aireka Scientific, HK). Fluorescence imaging was conducted utilizing a spinning-disk confocal microscope equipped with a Yokogawa CSU-W1 (Yokogawa Electric, Japan) scanner unit, which was integrated with a Nikon Ti2-E inverted microscope (Nikon Instruments Inc., US). Fluorophores were excited through the iLAS laser launcher (Gataca Systems, France) housing 405, 488, 561, and 642 nm laser lines. Image z-stacks (z-steps=0.3 μm) were acquired using a 40× water immersion objective (CFI Apo LWD 40XWI λS N.A. 1.15, Nikon) and captured with a sCMOS Camera (Prime 95B 22 mm, Teledyne Photometrics, US). The z-positions were precisely controlled via a piezo stage (PI PIFOC Z-stage, Physik Instrumente, US). The entire microscopy system was coordinated using the MetaMorph advanced acquisition software (Molecular Devices, USA).

## Analysis of actin alignment

Imaging of the various substrates resulted in multiple 3D time-lapse image stacks. To facilitate the downstream analysis of these image stacks, they were unwrapped to obtain a 2D representation of the monolayer. Unwrapping was done by first drawing the side profiles of the wave structures manually. A coordinate map, which detailed the correspondence of the coordinates in 3D space to the 2D plane, was generated from these side profiles. This was then used to unwrap the 3D image stack via spline interpolation.

Hill and valley regions of the unwrapped image were then extracted and separately quantified for alignment using the directionality function in FIJI with frequency binned into 90 bins. The data was then imported into the statistical programming software R for statistical analysis and plotting. 3 independent regions from 3 independent samples were analyzed for each dimension and time condition.

For the statistical tests a Two-way ANOVA was used, and the Tukey post hoc test was performed for the subsequent multiple comparisons.

## ACKNOWLEDGMENTS

We would like to thank the National University of Singapore (NUS) Mechanobiology Institute for the support, facilities and resources provided during this work.

## REFERENCES

1. D. W. Thompson, On growth and form (Cambridge Univ. Press, Cambridge, 1942).

2. D. Baptista, L. Teixeira, C. van Blitterswijk, S. Giselbrecht, R. Truckenmüller, Overlooked? underestimated? effects of substrate curvature on cell behavior. Trends in Biotechnology 37, 838–854 (2019). Doi: 10.1016/j.tibtech.2019.01.006

3. S. J. P. Callens, R. J. C. Uyttendaele, L. E. Fratila-Apachitei, A. A. Zadpoor, Substrate curvature as a cue to guide spatiotemporal cell and tissue organization. Biomaterials 232, 119739 (2020). Doi: 10.1016/j.biomaterials.2019.119739

4. C.-K. Huang, G. J. Paylaga, S. Bupphathong, K.-H. Lin, Spherical microwell arrays for studying single cells and microtissues in 3D confinement. Biofabrication 12, 025016 (2020). Doi: 10.1088/1758-5090/ab6eda

5. J.-Y. Lin, W.-J. Lin, W.-H. Hong, W.-C. Hung, S. H. Nowotarski, S. M. Gouveia, I. Cristo, K.-H. Lin, Morphology and organization of tissue cells in 3D microenvironment of monodisperse foam scaffolds. Soft Matter 7, 10010 (2011). Doi: 10.1039/C1SM05371J

6. P. Weiss, Experiments on cell and axon orientation in vitro: The role of colloidal exudates in tissue organization. Journal of Experimental Zoology 100, 353–386 (1945). Doi: 10.1002/jez.1401000305

7. Y. A. Rovensky, V. I. Samoilov, Morphogenetic response of cultured normal and transformed fibroblasts, and epitheliocytes, to a cylindrical substratum 16 surface: Possible role for the actin filament bundle pattern. Journal of Cell Science 107, 1255–1263 (1994). Doi: 0.1242/jcs.107.5.1255

8. C. M. Hwang, Y. Park, J. Y. Park, K. Lee, K. Sun, A. Khademhosseini, S. H. Lee, Controlled cellular orientation on PLGA microfibers with defined diameters. Biomedical Microdevices 11, 739–746 (2009). Doi: 10.1007/s10544-009-9287-7

9. S. M. Yu, B. Li, F. Amblard, S. Granick, Y. K. Cho, Adaptive architecture and mechanoresponse of epithelial cells on a torus. Biomaterials 265, 120420 (2021). Doi: 10.1016/j.biomaterials.2020.120420

10. V. Hosseini, P. Kollmannsberger, S. Ahadian, S. Ostrovidov, H. Kaji, V. Vogel, A. Khademhosseini, Fiber-Assisted Molding (FAM) of surfaces with tunable curvature to guide cell alignment and complex tissue architecture. Small 10, 4851–4857 (2014). Doi: 10.1002/smll.201400263

11. K. H. Song, S. J. Park, D. S. Kim, J. Doh, Sinusoidal wavy surfaces for curvature-guided migration of Tlymphocytes. Biomaterials 51, 151–160 (2015). Doi: 10.1016/j.biomaterials.2015.01.071

12. L. Pieuchot, J. Marteau, A. Guignandon, T. Dos Santos, I. Brigaud, P. Chauvy, T. Cloatre, A. Ponche, T. Petithory, P. Rougerie, M. Vassaux, J. Milan, N. T. Wakhloo, A. Spangenberg, M. Bigerelle, K. Anselme, Curvotaxis directs cell migration through cell-scale curvature landscapes. Nature Communications 9, 3995 (2018). Doi: 10.1038/s41467-018-06494-6

13. C.-K. Huang, X. Yong, D. T. She, C. T. Lim, Surface curvature and basal hydraulic stress induce spatial bias in cell extrusion. eLife 12:RP84921 (2023). Doi: 10.7554/eLife.84921.1

14. A. Jaiswal, C. K. Rastogi, S. Rani, G. P. Singh, S. Saxena, S. Shukla, Two decades of two-photon lithography: Materials science perspective for additive manufacturing of 2D/3D nano-microstructures. iScience 26, 106374 (2023). Doi: 10.1016/j.isci.2023.106374

15. J. Luo, M. Walker, Y. Xiao, H. Donnelly, M. J. Dalby, M. Salmeron-Sanchez, The influence of nanotopography on cell behaviour through interactions with the extracellular matrix – A review. Bioactive Materials 15, 145–159 (2022). Doi: 10.1016/j.bioactmat.2021.11.024

16. C.-K. Huang, A. Donald, Revealing the dependence of cell spreading kinetics on its spreading morphology using microcontact printed fibronectin patterns. Journal of The Royal Society Interface 12, 20141064–20141064 (2014). Doi: 10.1098/rsif.2014.1064

17. K. M. C. Leong, M. H. Nai, F. C. Cheong, C. T. Lim. Viscoelastic effects of silicone gels at the micro-and nanoscale. Procedia IUTAM 12, 20–30 (2015). Doi: 10.1016/j.piutam.2014.12.004

18. C. Moraes, J. M. Labuz, Y. Shao, J. Fu, S. Takayama, Supersoft lithography: candy-based fabrication of soft silicone microstructures. Lab Chip 15, 3760–5 (2015). Doi: 10.1039/c5lc00722d

